# CRF_1_ receptor-dependent increases in irritability-like behavior during abstinence from chronic intermittent ethanol vapor exposure

**DOI:** 10.1101/159871

**Authors:** Adam Kimbrough, Giordano de Guglielmo, Jenni Kononoff, Marsida Kallupi, Eric P. Zorrilla, Olivier George

**Author notes:** Correspondence should be addressed to, Dr. Olivier George, ^1^Department of Neuroscience, The Scripps Research Institute, 10550 North 11 Torrey Pines Road, SP30-2400, La Jolla, CA 92037.

## Abstract

**Background:** In humans, emotional and physical signs of withdrawal from ethanol are commonly seen. Many of these symptoms, including anxiety-like and depression-like behavior, have been characterized in animal models of ethanol dependence. One issue with several current behavioral tests measuring withdrawal in animal models is they are often not repeatable within subjects over time. Additionally, irritability, one of the most common symptoms of ethanol withdrawal in humans, has not been well characterized in animal models. The corticotropin-releasing factor (CRF)-CRF_1_ receptor system has been suggested to be critical for the emergence of anxiety-like behavior in ethanol dependence, but the role of this system in irritability-like behavior has not been characterized.

**Methods:** The present study compared the effects of chronic intermittent ethanol vapor exposure (CIE)-induced ethanol dependence on irritability-like behavior in rats using the bottle-brush test during acute withdrawal and protracted abstinence. Rats were trained to self-administer ethanol in operant chambers and then either left in a nondependent state or made dependent via CIE. Naive, nondependent, and dependent rats were tested for irritability-like behavior in the bottle-brush test 8 h and 2 weeks into abstinence from ethanol. A separate cohort of dependent rats was used to examine the effect of the specific CRF_1_ receptor antagonist R121919 on irritability-like behavior.

**Results:** Dependent rats exhibited escalated ethanol intake compared with their own pre-CIE baseline and nondependent rats. At both time-points of abstinence, ethanol-dependent rats exhibited increased aggressive-like responses compared with naive and nondependent rats. R121919 blocked the increased irritability-like behavior in dependent rats.

**Conclusions:** The effect of R121919 to block increased irritability-like behavior suggests that CRF plays an important role in this behavior, similar to other negative emotional withdrawal symptoms. Quantifying and understanding the molecular basis of irritability-like behavior may yield new insights into withdrawal from ethanol and other drugs of abuse.

## 1. Introduction

Alcoholism is a chronic relapsing disorder associated with compulsive drinking, the loss of control over intake, and the emergence of a negative emotional state during abstinence (Koob et al., 2004, Koob and Volkow, 2010). In humans, the emotional and physical signs of ethanol withdrawal include anxiety, irritability, mood swings, insomnia, tremors, convulsions, higher blood pressure, accelerated pulse, accelerated breathing, accelerated heart rate, dehydration, and delirium tremens (Schuckit et al., 1995, Finn and Crabbe, 1997). The emotional symptoms of ethanol dependence, including anxiety-like behavior and depression-like behavior, can be modeled in animals during ethanol dependence (Pleil et al., 2015, Marcinkiewcz et al., 2015, Thorsell et al., 2007, Varlinskaya et al., 2017, Kallupi et al., 2014, Gilpin et al., 2012, Gilpin et al., 2015, Heilig et al., 2010, Kliethermes et al., 2004, Pandey et al., 2015, Pandey et al., 2003, Valdez et al., 2004, Valdez et al., 2002a, Ehlers et al., 2013, Vetreno et al., 2016, Rylkova et al., 2009, McClintick and Grant, 2016, Egli et al., 2012, Buck et al., 2014). An important challenge for preclinical researchers is the fact that these tests are usually not repeatable within subjects over time, which prevents conducting reliable longitudinal studies. Moreover, increased irritability is a key negative emotional symptom that has been largely neglected.

Irritability is one of the most common withdrawal symptoms in humans (Lubman et al., 1983) and has been anecdotally noted during withdrawal in animal models but shown to be difficult to quantify experimentally (Frye and Ellis, 1977, Riihioja et al., 1997, Riihioja et al., 1999, Woldbye et al., 2002, Becker, 2000). Thus, it is critical to characterize irritability-like behavior during withdrawal and protracted abstinence from ethanol in dependent rats.

A commonly used method to study ethanol dependence and withdrawal is the chronic intermittent ethanol (CIE) vapor exposure model (Kissler et al., 2014, Gilpin et al., 2008c, Gilpin et al., 2008b, Staples et al., 2015, Leao et al., 2015, de Guglielmo et al., 2016, Kimbrough et al., 2017). Rats that are made dependent with CIE exhibit clinically-relevant blood ethanol levels (BELs; 150-250 mg%), an increase in ethanol drinking when tested during early and protracted withdrawal, and compulsive-like ethanol drinking (i.e., responding despite adverse consequences; (Rogers et al., 1979, Roberts et al., 1996, Roberts et al., 2000, O’Dell et al., 2004, Schulteis et al., 1995, Vendruscolo et al., 2012, Seif et al., 2013, Leao et al., 2015, Kimbrough et al., 2017). Ethanol dependence that is induced by ethanol vapor results in withdrawal symptoms during both acute withdrawal (somatic and motivational) and protracted abstinence (motivational; (Macey et al., 1996, Sommer et al., 2008, Schulteis et al., 1995, Williams et al., 2012, Kallupi et al., 2014, Valdez et al., 2002b, Zhao et al., 2007, Vendruscolo and Roberts, 2014, de Guglielmo et al., 2017), but the effects of abstinence from CIE on irritability-like behavior during withdrawal and after protracted abstinence has not yet been reported.

Converging lines of evidence suggest that recruitment of the brain corticotropin-releasing factor (CRF)-CRF_1_ receptor system during withdrawal from CIE is critical for the emergence of anxiety-like behavior (Valdez et al., 2002a, Sabino et al., 2006, Finn et al., 2007, Gilpin et al., 2008b, Richardson et al., 2008b, Richardson et al., 2008a), but the role of the CRF-CRF_1_ system in irritability-like behavior during withdrawal remains to be demonstrated. In the present study, we hypothesized that (1) abstinence from ethanol after the escalation of ethanol drinking using the CIE model would increase irritability-like behavior in the bottle-brush test 8 h into withdrawal and after 2 weeks of protracted abstinence (Riittinen et al., 1986, Lagerspetz and Portin, 1968) and (2) the CRF_1_ receptor antagonist R121919 would reduce irritability-like behavior during withdrawal.

## Materials and Methods

### Animals

Adult male Wistar rats (Charles River), weighing 250-300 g at the beginning of the experiments, were used. The rats were group housed, two per cage, in a temperature-controlled (22°C) vivarium on a 12 h/12 h light/dark cycle (lights on at 10:00 PM) with *ad libitum* access to food and water. All of the procedures were conducted in strict adherence to the National Institutes of Health *Guide for the Care and Use of Laboratory Animals* and approved by The Scripps Research Institute Institutional Animal Care and Use Committee.

### Operant ethanol self-administration

Two cohorts of rats were used in this experiment. One cohort of rats (*n* = 24) and an additional cohort of rats (*n* = 16) were trained to self-administer ethanol. Both cohorts self-administered 10% (w/v) ethanol during daily sessions in standard operant conditioning chambers (Med Associates) until stable responding was maintained as previously described (de Guglielmo et al., 2016, Leao et al., 2015, Kimbrough et al., 2017). The rats were first subjected to an overnight session in the operant chambers with access to one lever (front lever) that delivered water on a fixed-ratio 1 (FR1) schedule (i.e., each operant response was reinforced with 0.1 ml of the solution). Food was available *ad libitum* during this training period. After 1 day off, the rats were subjected to a 3 h session (FR1) for 1 day, a 2 h session (FR1) the next day, and a 1 h session (FR1) the next day, with one lever delivering ethanol (front lever, 0.1 ml). All of the subsequent sessions lasted 30 min, and two levers were available (front lever: ethanol; back lever: water) until stable levels of intake were reached. Upon completion of this procedure, the animals were allowed to self-administer a 10% (w/v) ethanol solution and water on an FR1 schedule of reinforcement. The animals in the first cohort were then divided into two groups (*n* = 12 dependent, *n* = 12 nondependent). The dependent group underwent the CIE protocol, and the nondependent group was left undisturbed in the vivarium. All of the rats in the second cohort underwent the CIE protocol to become ethanol-dependent.

### Chronic intermittent ethanol vapor

The rats in the dependent groups were made ethanol-dependent by the CIE vapor procedure as previously described (O’Dell et al., 2004, Gilpin et al., 2008a). The rats underwent repeated daily cycles of 14 h vapor ON (blood ethanol levels during vapor exposure ranged from 150 to 250 mg%) and 10 h vapor OFF, during which behavioral testing occurred (i.e., 6-8 h after the vapor was turned OFF), when brain and BELs are negligible (Gilpin et al., 2009). After 4-6 weeks of vapor exposure, the rats resumed operant self-administration sessions during withdrawal to test for the escalation of ethanol intake.

### Blood ethanol measurements

Tail blood was collected and used to determine blood ethanol levels (BELs) using an oxygen-rate ethanol analyzer (Analox Instruments, Stourbridge, UK).

### Irritability-like behavior

To test irritability-like behavior during ethanol withdrawal and protracted abstinence, we used the bottle-brush test, based on the methods of Riittinen et al. (1986) and Lagerspetz and Portin (1968) and modified slightly for rats. Irritability-like behavior was tested following the escalation of ethanol intake in the cohorts of dependent and nondependent rats and in age-matched ethanol-naive rats (*n* = 10). Testing occurred after 8 h of withdrawal or 2 weeks of protracted abstinence from CIE (in the dependent group). Irritability-like behavior was examined by measuring aggressive and defensive responses during the bottle-brush test.

Irritability-like behavior testing was performed in the middle half of the dark cycle under dark conditions with a red light for the observers. The sessions were conducted in a randomized order for each animal. Testing consisted of 10 trials per rat in plastic cages (10.5 in × 19 in × 8 in; Ancare, Bellmore, NY, USA) with fresh bedding. During each trial, the rat started at the back of the cage. A bottle-brush was rotated toward the animal’s whiskers (from the front of the cage) by a treatment-naive experimenter. The brush was rotated around the whiskers of the rat for approximately 1 s. The brush was then rotated back to the front of the cage where it was allowed to hang vertically for approximately 2 s, during which behavioral responses were recorded. A 10-s intertrial interval was used. Three observers who were blind to treatment scored the behaviors in real time.

For each rat, separate sums of aggressive and defensive responses across all trials were determined for each observer. Aggressive and defensive response scores for each rat were then calculated by averaging the observers’ sums. This was then used to calculate a group mean and SEM.

The following were scored as aggressive responses: smelling the target, biting the target (during the initial phase of rotating the brush forward and back to the starting position), boxing the target, following the target, exploring the target (using paws or mouth to manipulate the brush without biting or boxing), mounting the target, and delayed biting (during the 2 s that the brush hung at the starting position). The following were scored as defensive responses: escaping from the target, digging, burying, jumping, climbing, vocalization, freezing, and grooming. Grooming and digging were additionally recorded during the 10-s intertrial intervals.

### Effects of CRF antagonist on irritability-like behavior

The second cohort of rats that was made dependent on ethanol via CIE exposure was tested for irritability-like behavior 8 h into withdrawal. Thirty minutes before testing, the rats were injected i.p. with either vehicle or the selective CRF_1_ receptor antagonist R121919 (10 mg/kg; synthesized by Dr. Kenner Rice at the Drug Design and Synthesis Section, Chemical Biology Research Branch, National Institute on Drug Abuse, National Institutes of Health, Bethesda, MD). Both vehicle and R121919 were administered in a solution that contained 5% dimethylsulfoxide, 5% Emulphor, and 90% distilled water. Following the injection of vehicle or R121919, the rats underwent the bottle-brush test as described above.

### Statistical analysis

The results are expressed as mean ± SEM. For the cohorts of dependent and nondependent rats, the last 3 days of ethanol intake before vapor exposure were averaged to obtain the pre-vapor baseline intake. The last 3 days of ethanol intake before bottle-brush testing were averaged to obtain the post-vapor (escalated) intake. The data were analyzed using repeated-measures analysis of variance (ANOVA), with group (nondependent and dependent) as the between-subjects factor and day (baseline intake and escalated intake) as the within-subjects factor. The data were also analyzed by week using repeated-measures ANOVA, with group (nondependent and dependent) as the between-subjects factor and week as the within-subjects factor. Behavioral data for the second cohort were analyzed using a one-way ANOVA to compare the average of the last 3 days of ethanol intake before vapor exposure (pre-vapor baseline intake) and the average of the last 3 days of ethanol intake before bottle-brush testing (post-vapor escalated intake). For the bottle-brush test, each time point (8 h of withdrawal and 2 weeks of protracted abstinence) was examined using a one-way ANOVA. The ANOVAs were followed by Fisher’s Least Significant Difference (LSD) *post hoc* test when appropriate. To evaluate the effects of R121919 on behavior in the bottle-brush test, *t*-tests were performed between the two groups. Differences were considered significant at *p* < 0.05. All of the data were analyzed using Statistica 13 software (StatSoft, Palo Alto, USA).

## Results

### Blood ethanol levels

Blood ethanol levels were measured during CIE. In dependent rats, BELs were maintained between 150 and 250 mg/100 ml with no differences found between groups (data not shown).

### Operant ethanol self-administration during CIE exposure

For operant ethanol self-administration, the mixed factorial ANOVA, with group (nondependent and dependent) as the between-subjects factor and day (baseline intake and escalated intake) as the within-subjects factor, revealed a significant day × group interaction (*F*_1,22_ = 7.5, *p* < 0.05) and significant effects of day (*F*_1,22_ = 5.4, *p* <0.05) and group (*F*_1,22_ = 4.1, *p* = 0.05). Fisher’s LSD *post hoc* test revealed that both dependent and nondependent rats reached a stable baseline of responding for ethanol during training (36.8 ± 5.3 lever presses in dependent rats *vs*. 36.1 ± 4.7 lever presses in nondependent rats), with no significant difference between groups. After CIE exposure, dependent rats significantly escalated the number of lever presses for ethanol compared with nondependent rats (61.9 ± 7.7 lever presses in dependent rats *vs*. 34.0 ± 6.3 lever presses in nondependent rats; *p* < 0.05; Fig 1A) and compared with their own baseline responding (61.9 ± 7.7 lever presses in dependent CIE rats *vs*. 36.8 ± 5.3 lever presses in dependent baseline rats; *p* < 0.05; Fig. 1A).

**Figure 1.**
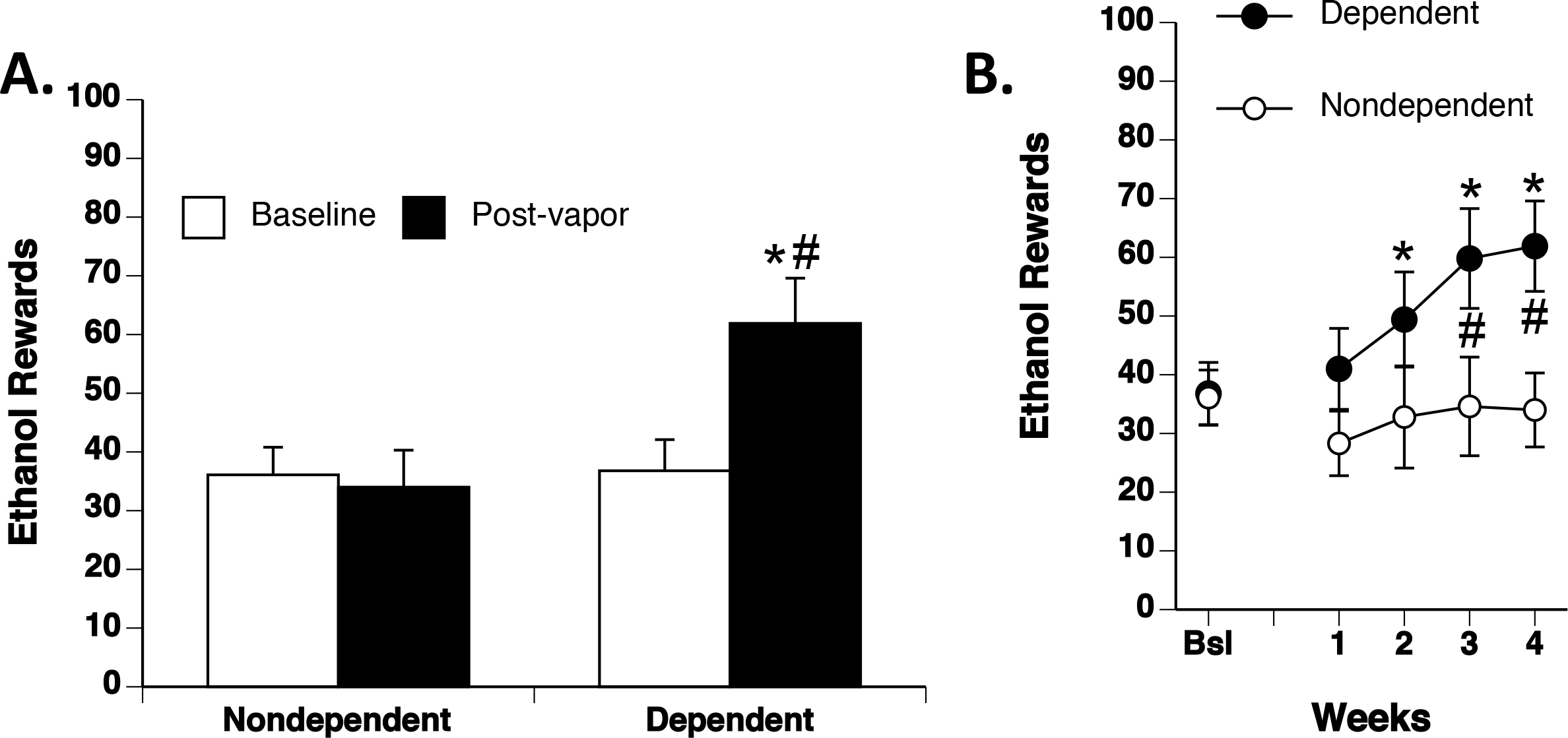
Escalation of operant ethanol self-administration after CIE. Rats were trained to operantly self-administer ethanol until they reached a stable baseline. Dependent rats were made dependent via chronic intermittent ethanol vapor. Dependent and nondependent rats were then tested daily for the escalation of operant ethanol self-administration. **(A)** Average of the last 3 days of pre-vapor (baseline) and post-vapor (escalation) prior to testing irritability-like behavior. Dependent rats significantly escalated their ethanol intake relative to baseline (white bar) after CIE (post vapor; black bar). Dependent rats also exhibited significantly higher ethanol intake after CIE compared with nondependent rats (post vapor; black bar). Nondependent rats did not significantly escalate their ethanol intake relative to baseline (white bar). **(B)** Average ethanol intake per operant session during each week of CIE exposure. Dependent rats (black circles) significantly escalated their ethanol intake relative to their own baseline at weeks 2-4. Dependent rats significantly escalated their ethanol intake compared with nondependent rats (white circles) at weeks 3 and 4. \**p* < 0.05, dependent post-vapor each week *vs*. dependent baseline intake; *^#^p* < 0.05, dependent post-vapor each week *vs*. nondependent post-vapor each week.

When we analyzed the data according to weeks of ethanol self administration, the repeated-measures ANOVA, with group (nondependent and dependent) as the between-subjects factor and week as the within-subjects factor, revealed a significant week × group interaction (*F*_4,88_ = 3.9, *p* < 0.05) and a significant effect of week (*F*_4,88_ = 4.8, *p* < 0.005) but no effect of group (*F*_1,22_ = 3.6, *p* = 0.07). Fisher’s LSD *post hoc* test revealed that dependent rats presented a significant increase in ethanol intake relative to their own baseline at weeks 2-4 of CIE exposure. Dependent rats also exhibited a significant increase in ethanol intake compared with nondependent rats at weeks 3 and 4 (Fig. 1B).

### Irritability-like behavior at 8 h withdrawal and 2 weeks abstinence from CIE

When tested 8 h into withdrawal from ethanol vapor, there was a significant effect of group on aggressive responses (*F*_2,31_ = 11.8, *p* < 0.0005). Fisher’s LSD *post hoc* test showed that ethanol-dependent rats had a significantly higher number of aggressive responses (26.3 ± 2.8) compared with both nondependent rats (14.4 ± 2.6) and naive rats (9.6 ± 1.6; Fig. 2A). No significant difference in defensive responses was found between groups (Fig. 2B).

**Figure 2.**
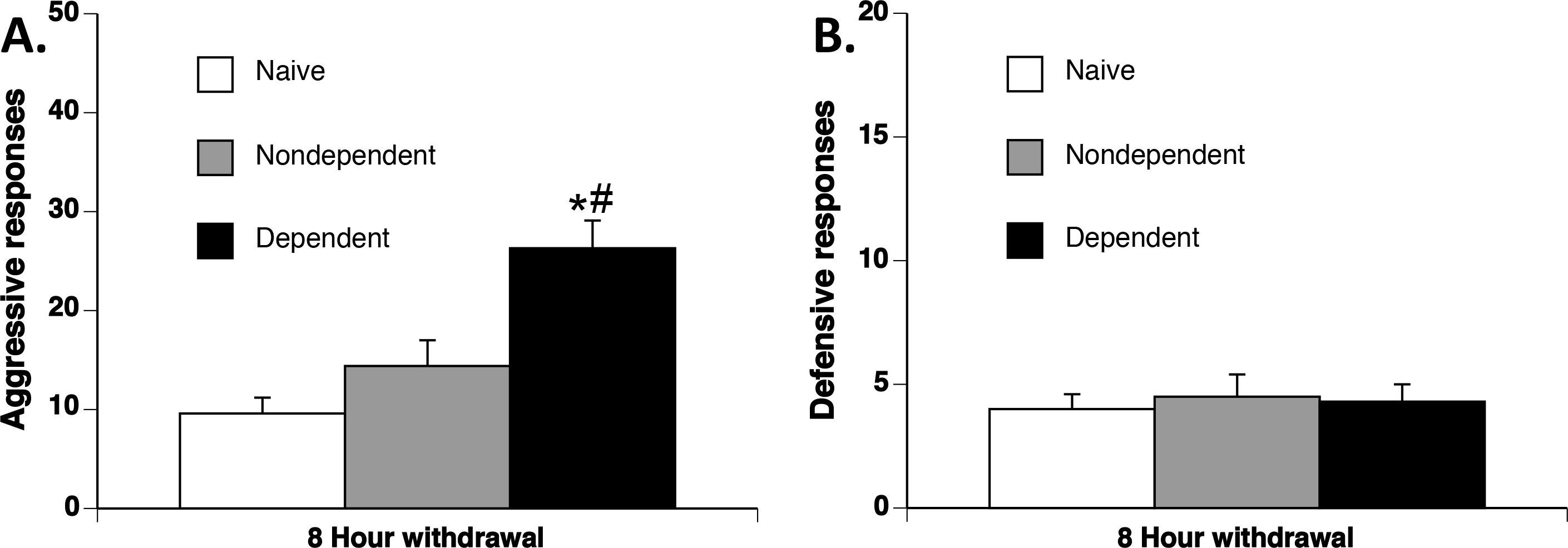
Irritability-like behavior in ethanol-dependent rats 8 h into withdrawal from ethanol vapor. Rats underwent the bottle-brush test to assess aggressive and defensive responses. **(A)** Aggressive responses 8 h into withdrawal. Dependent rats (black bar) exhibited a significant increase in the number of aggressive responses over the course of 10 trials compared with nondependent rats (gray bar) and naive rats (white bar). **(B)** Defensive responses 8 h into withdrawal. No significant differences in the number of defensive responses were found between groups. ^*^*p* < 0.05, dependent *vs*. naive; ^#^*p* < 0.05, dependent *vs*. nondependent.

When tested 2 weeks into protracted abstinence from ethanol vapor, there was a significant effect of group on aggressive responses (*F*_2,31_ = 6.7, *p* < 0.005). Fisher’s LSD *post hoc* test showed that ethanol-dependent rats had a significantly higher number of aggressive responses (27.2 ± 2.7) compared with both nondependent rats (18 ± 2.0) and naive rats (14.8 ± 2.7; Fig. 3A). No significant difference in defensive responses was found between groups (Fig. 3B).

**Figure 3.**
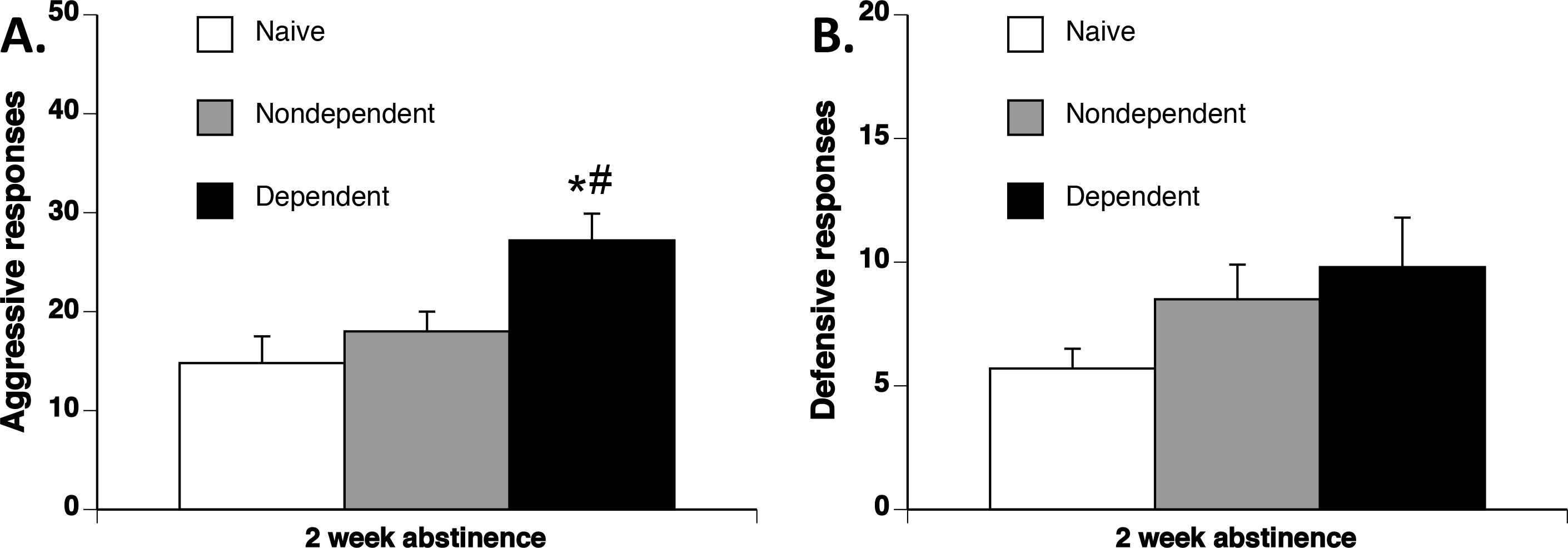
Irritability-like behavior following 2 weeks of protracted abstinence from ethanol vapor in dependent rats. Rats underwent the bottle-brush test to assess aggressive and defensive responses. **(A)** Aggressive responses after 2 weeks of protracted abstinence. Dependent rats (black bar) exhibited a significant increase in the number of aggressive responses over the course of 10 trials compared with nondependent rats (gray bar) and naive rats (white bar). **(B)** Defensive responses after 2 weeks of protracted abstinence. No significant differences in the number of defensive responses were found between groups. \**p* < 0.05, dependent *vs*. naive; *^#^p* < 0.05, dependent *vs*. nondependent.

### Effects of CRF antagonist on irritability-like behavior

Operant ethanol self-administration for the vehicle (baseline: 14.8 ± 4.2 lever presses; escalated: 46.8 ± 8.3 lever presses) and R121919 (baseline: 15.8 ± 4.5 lever presses; escalated: 47.5 ± 7.5 lever presses) groups did not differ at either baseline or during escalation. The data from both groups were combined, and further analyses were performed to compare baseline *vs*. escalated intake in all dependent rats using a one-way repeated-measures ANOVA, with day (baseline intake and escalated intake) as the within-subjects factor. There was a significant effect of day on responding for ethanol (*F*_1,15_ = 22.1, *p* < 0.0005). Fisher’s LSD *post hoc* test revealed that ethanol-dependent rats presented significantly higher ethanol intake during escalation compared with their own baseline (15.3 ± 3.0 lever presses at baseline *vs*. 47.1 ± 5.4 lever presses during escalation; Fig. 4A).

**Figure 4.**
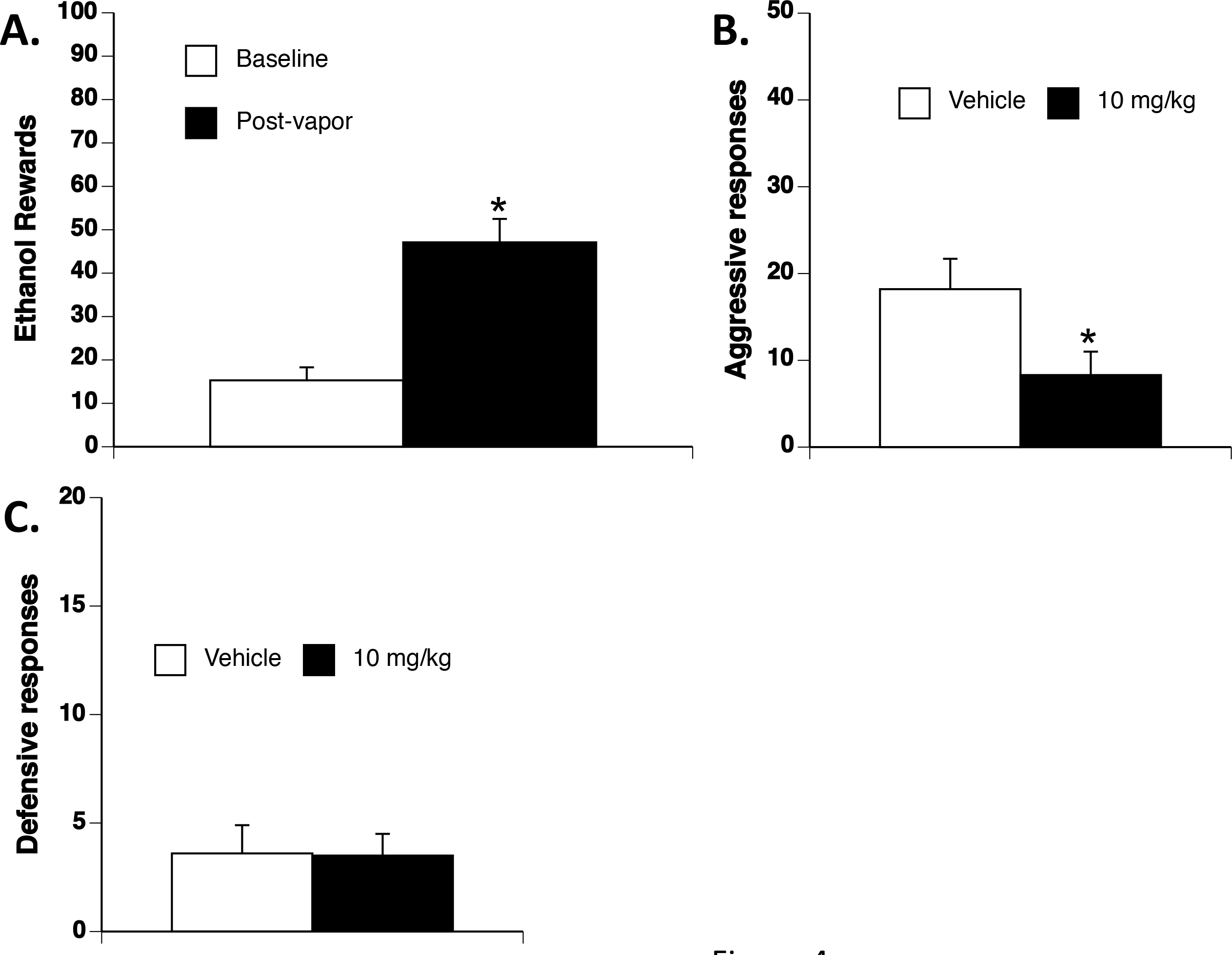
Escalation of ethanol intake and irritability-like behavior in ethanol-dependent rats 8 h into withdrawal from ethanol vapor. Rats were given CIE exposure to escalate ethanol intake relative to baseline. Escalated rats were then treated with vehicle or the CRF_1_ receptor antagonist R121919 (10 mg/kg) 30 min prior to testing. The rats then underwent the bottle-brush test to assess aggressive and defensive responses. **(A)** Average of the last 3 days pre-vapor (baseline) and post-vapor (escalation) prior to testing irritability-like behavior. Dependent rats significantly escalated their ethanol intake relative to baseline (white bar) after CIE (post-vapor; black bar). **(B)** Aggressive responses. R121919-treated rats (black bar) exhibited a significant reduction of the number of aggressive responses over the course of 10 trials compared with vehicle-treated rats (white bar). **(C)** Defensive responses. No significant differences in the number of defensive responses were found between groups. \**p* < 0.05, R121919 *vs*. vehicle.

When ethanol-dependent rats were tested for irritability-like behavior in the bottle-brush test during acute withdrawal (after injection of vehicle or R121919), there was a significant effect of group (vehicle vs. 10 mg/kg R121919; *t*_14_ = 2.2, *p* < 0.05). R121919 significantly decreased aggressive responses (8.3 ± 2.7) compared with vehicle-treated rats (18.2 ± 3.5; Fig. 4B). No significant difference in defensive responses was found between groups (Fig. 4C).

## Discussion

The present study found that rats that were made dependent on ethanol via CIE exhibited an increase in irritability-like behavior in the bottle-brush test 8 h into withdrawal and 2 weeks into protracted abstinence. The specific CRF_1_ receptor antagonist R121919, at a dose that is known to block central CRF_1_ receptors, decreased the withdrawal-induced increase in irritability-like behavior.

Similar to other studies (Gilpin et al., 2008c, Gilpin et al., 2008b, Leao et al., 2015, Williams et al., 2012, O’Dell et al., 2004, Walker et al., 2008), rats that were made dependent on ethanol via CIE significantly escalated their ethanol intake. Interestingly, we found that ethanol-dependent rats exhibited an increase in irritability-like behavior in the bottle-brush test 8 h into withdrawal and 2 weeks into protracted abstinence from ethanol vapor compared with both nondependent and naive rats. R121919 blocked the increase in irritability-like behavior 8 h into withdrawal in dependent rats. This effect was observed only for aggressive responses and not for defensive responses, indicating a behaviorally-specific effect.

Aggressive irritability-like behavior has not been previously tested to measure negative emotional states during ethanol withdrawal in the rat CIE model. Animal models of irritability have been difficult to characterize, and reports are often limited to anecdotes or have used less specific or quantitative assessments, such as reactivity to handling (Frye and Ellis, 1977, Riihioja et al., 1997, Riihioja et al., 1999, Woldbye et al., 2002, Becker, 2000). An increase in irritability-like behavior 8 h into withdrawal from ethanol is consistent with results for other withdrawal symptoms.

Somatic withdrawal symptoms have been well characterized during withdrawal from ethanol vapor. Somatic withdrawal symptoms, such as tremors, abnormal gait, and tail stiffness, are often observed during acute withdrawal from ethanol vapor. These changes can be seen as early as 2 h into withdrawal and last up to 72 h (Macey et al., 1996, de Guglielmo et al., 2016, de Guglielmo et al., 2015). The motivational symptoms of withdrawal have been found to last much longer (> 2 weeks) into protracted abstinence. Increases in anxiety-like behavior in the elevated plus maze and depressive-like behavior (e.g., immobility) in the forced swim test have been reported in ethanol-dependent rats and mice during acute withdrawal from ethanol vapor and after protracted abstinence (Kallupi et al., 2014, Valdez et al., 2002b, Zhao et al., 2007, Williams et al., 2012, Marcinkiewcz et al., 2015, Shibasaki et al., 2012, Walker et al., 2010). Impairments in brain reward function, reflected by elevations of intracranial self-stimulation (ICSS) thresholds, have also been observed during acute withdrawal from ethanol vapor in dependent rats (Schulteis et al., 1995). Perhaps the closest measures of irritability that have been reported during withdrawal include 22 kHz ultrasonic vocalizations and measures of reactivity to touch or handling (including vocalizations; (Frye and Ellis, 1977, Riihioja et al., 1997, Riihioja et al., 1999, Woldbye et al., 2002, Becker, 2000, Akunne and Soliman, 1988). Ultrasonic vocalizations are thought to represent distress in animals (Becker, 2000) and greatly increase during withdrawal from ethanol vapor and an ethanol liquid diet (Williams et al., 2012, Knapp et al., 1998, Buck et al., 2014). Neither of these measures, however, recapitulates the specific increase in aggressive responses that were observed in the present study, which might more specifically model the quick-to-anger, grouchiness, and hostility that is seen during ethanol withdrawal in humans.

In the present study, we observed an increase in irritability-like behavior after 2 weeks of protracted abstinence, which is consistent with motivational measures of ethanol withdrawal that have been shown to persist into protracted abstinence (2-8 weeks) from ethanol vapor (Roberts et al., 2000). Rats that underwent 4-6 weeks of abstinence from ethanol exhibited an increase in anxiety-like behavior and a decrease in the time spent on the open arms in the elevated plus maze (Valdez et al., 2002b, Zhao et al., 2007). However, an important advantage of using the bottle-brush test over the elevated plus maze test is that the former can be repeated in the same individuals. Our results revealed a consistent increase in irritability-like behavior during both acute and protracted abstinence in the same subjects.

Altogether, the present results suggest that irritability-like behavior may be a clinically relevant measure of a negative emotional state that may model the anxiety, irritability, and mood disturbances that can persist long after detoxification in ethanol-dependent humans (Brady et al., 2002, Martinotti et al., 2007, Miyata et al., 2008, Haney, 2005). Irritable aggression may be a prognostically relevant motivational withdrawal symptom that predicts an increase in craving (Chiang et al., 2002) and more relapse episodes in ethanol-dependent males during abstinence (Baars et al., 2013).

CRF-CRF_1_ receptor signaling has been shown to be critically involved in ethanol withdrawal-related behaviors (Zorrilla et al., 2013). CRF blockade with various receptor antagonists has been shown to reduce ethanol intake in dependent rats (Heilig and Koob, 2007, Funk et al., 2007, Valdez et al., 2002b, Funk et al., 2006) and anxiety-like behavior in dependent rats (Heilig and Koob, 2007, Baldwin et al., 1991, Rassnick et al., 1993, Valdez et al., 2003). Irritability-like behavior also appears to be inhibited by CRF_1_ receptor blockade, suggesting that irritability- and anxiety-like behavior may share common neurocircuitry. Combining the bottle-brush test with other reliable tests may help identify unique *vs*. overlapping neural substrates and the pathophysiological relevance of different motivational symptoms of withdrawal.

In summary, the present study found increases in irritable aggression-like behavior in ethanol-dependent rats during acute withdrawal and protracted abstinence. CRF_1_ antagonism specifically blocked the increase in irritability-like behavior in ethanol-dependent rats. The bottle-brush test may represent a clinically relevant, reliable, and reproducible method to investigate the neurobiological mechanisms that underlie the emergence of negative emotional states during abstinence and may facilitate drug discovery by providing a high-throughput within-subjects paradigm for the preclinical screening of novel medications for the treatment of symptoms of alcohol use disorders.

## Acknowledgements

The authors acknowledge expert technical assistance by Maury Cole and editorial assistance by Michael Arends.

## References

Akunne, H. C. & Soliman, K. F. 1988. Hyperglycemic suppression of morphine withdrawal signs in the rat. Psychopharmacology (Berl), 96, 1–6.

Baars, M. Y., Muller, M. J., Gallhofer, B. & Netter, P. 2013. Relapse (number of detoxifications) in abstinent male alcohol-dependent patients as related to personality traits and types of tolerance to frustration. Neuropsychobiology, 67, 241–248.

Baldwin, H. A., Rassnick, S., Rivier, J., Koob, G. F. & Britton, K. T. 1991. CRF antagonist reverses the “anxiogenic” response to ethanol withdrawal in the rat. Psychopharmacology (Berl), 103, 227–232.

Becker, H. C. 2000. Animal models of alcohol withdrawal. Alcohol Res Health, 24, 105–113.

Brady, K. T., Myrick, H., Henderson, S. & Coffey, S. F. 2002. The use of divalproex in alcohol relapse prevention: a pilot study. Drug Alcohol Depend, 67, 323–30.

Buck, C. L., Malavar, J. C., George, O., Koob, G. F. & Vendruscolo, L. F. 2014. Anticipatory 50 kHz ultrasonic vocalizations are associated with escalated alcohol intake in dependent rats. Behav Brain Res, 271, 171–176.

Chiang, S. S., Schuetz, C. G. & Soyka, M. 2002. Effects of irritability on craving before and after cue exposure in abstinent alcoholic inpatients: experimental data on subjective response and heart rate. Neuropsychobiology, 46, 150–60.

De Guglielmo, G., Crawford, E., Kim, S., Vendruscolo, L. F., Hope, B. T., Brennan, M., Cole, M., Koob, G. F. & George, O.. 2016. Recruitment of a Neuronal Ensemble in the Central Nucleus of the Amygdala Is Required for Alcohol Dependence. J Neurosci, 36, 9446–53.

De Guglielmo, G., Kallupi, M., Cole, M. D. & George, O. 2017. Voluntary induction and maintenance of alcohol dependence in rats using alcohol vapor self-administration. Psychopharmacology (Berl), 234, 2009–2018.

De Guglielmo, G., Martin-Fardon, R., Teshima, K., Ciccocioppo, R. & Weiss, F. 2015. MT-7716, a potent NOP receptor agonist, preferentially reduces ethanol seeking and reinforcement in post-dependent rats. Addict Biol, 20, 643–651.

Egli, M., Koob, G. F. & Edwards, S. 2012. Alcohol dependence as a chronic pain disorder. Neurosci Biobehav Rev, 36, 2179–2192.

Ehlers, C. L., Liu, W., Wills, D. N. & Crews, F. T. 2013. Periadolescent ethanol vapor exposure persistently reduces measures of hippocampal neurogenesis that are associated with behavioral outcomes in adulthood. Neuroscience, 244, 1–15.

Finn, D. A. & Crabbe, J. C. 1997. Exploring alcohol withdrawal syndrome. Alcohol Health Res World, 21, 149–156.

Finn, D. A., Snelling, C., Fretwell, A. M., Tanchuck, M. A., Underwood, L., Cole, M., Crabbe, J. C. & Roberts, A. J. 2007. Increased drinking during withdrawal from intermittent ethanol exposure is blocked by the CRF receptor antagonist D-Phe-CRF(12-41). Alcohol Clin Exp Res, 31, 939–949.

Frye, G. D. & Ellis, F. W. 1977. Effects of 6-hydroxydopamine or 5,7-dihydroxy-tryptamine on the development of physical dependence on ethanol. Drug Alcohol Depend, 2, 349–359.

Funk, C. K., O’Dell, L. E., Crawford, E. F. & Koob, G. F. 2006. Corticotropin-releasing factor within the central nucleus of the amygdala mediates enhanced ethanol self-administration in withdrawn, ethanol-dependent rats. J Neurosci, 26, 11324–11332.

Funk, C. K., Zorrilla, E. P., Lee, M. J., Rice, K. C. & Koob, G. F. 2007. Corticotropin-releasing factor 1 antagonists selectively reduce ethanol self-administration in ethanol-dependent rats. Biol Psychiatry, 61, 78–86.

Gilpin, N. W., Herman, M. A. & Roberto, M. 2015. The central amygdala as an integrative hub for anxiety and alcohol use disorders. Biol Psychiatry, 77, 859–869.

Gilpin, N. W., Karanikas, C. A. & Richardson, H. N. 2012. Adolescent binge drinking leads to changes in alcohol drinking, anxiety, and amygdalar corticotropin releasing factor cells in adulthood in male rats. PLoS One, 7, e31466.

Gilpin, N. W., Misra, K. & Koob, G. F. 2008a. Neuropeptide Y in the central nucleus of the amygdala suppresses dependence-induced increases in alcohol drinking. Pharmacol Biochem Behav, 90, 475–480.

Gilpin, N. W., Richardson, H. N. & Koob, G. F. 2008b. Effects of CRF1-receptor and opioid-receptor antagonists on dependence-induced increases in alcohol drinking by alcohol-preferring (P) rats. Alcohol Clin Exp Res, 32, 1535–42.

Gilpin, N. W., Richardson, H. N., Lumeng, L. & Koob, G. F. 2008c. Dependence-induced alcohol drinking by alcohol-preferring (P) rats and outbred Wistar rats. Alcohol Clin Exp Res, 32, 1688–96.

Gilpin, N. W., Smith, A. D., Cole, M., Weiss, F., Koob, G. F. & Richardson, H. N. 2009. Operant behavior and alcohol levels in blood and brain of alcohol-dependent rats. Alcohol Clin Exp Res, 33, 2113–2123.

Haney, M. 2005. The marijuana withdrawal syndrome: diagnosis and treatment. Curr Psychiatry Rep, 7, 360–366.

Heilig, M., Egli, M., Crabbe, J. C. & Becker, H. C. 2010. Acute withdrawal, protracted abstinence and negative affect in alcoholism: are they linked? Addict Biol, 15, 169–184.

Heilig, M. & Koob, G. F. 2007. A key role for corticotropin-releasing factor in alcohol dependence. Trends Neurosci, 30, 399–406.

Kallupi, M., Vendruscolo, L. F., Carmichael, C. Y., George, O., Koob, G. F. & Gilpin, N. W. 2014. Neuropeptide YY(2)R blockade in the central amygdala reduces anxiety-like behavior but not alcohol drinking in alcohol-dependent rats. Addict Biol, 19, 755–757.

Kimbrough, A., Kim, S., Cole, M., Brennan, M. & George, O. 2017. Intermittent Access to Ethanol Drinking Facilitates the Transition to Excessive Drinking After Chronic Intermittent Ethanol Vapor Exposure. Alcoholism: Clinical and Experimental Research, In Press.

Kissler, J. L., Sirohi, S., Reis, D. J., Jansen, H. T., Quock, R. M., Smith, D. G. & Walker, B. M. 2014. The one-two punch of alcoholism: role of central amygdala dynorphins/kappa-opioid receptors. Biol Psychiatry, 75, 774–782.

Kliethermes, C. L., Cronise, K. & Crabbe, J. C. 2004. Anxiety-like behavior in mice in two apparatuses during withdrawal from chronic ethanol vapor inhalation. Alcohol Clin Exp Res, 28, 1012–9.

Knapp, D. J., Duncan, G. E., Crews, F. T. & Breese, G. R. 1998. Induction of Fos-like proteins and ultrasonic vocalizations during ethanol withdrawal: further evidence for withdrawal-induced anxiety. Alcohol Clin Exp Res, 22, 481–493.

Koob, G. F., Ahmed, S. H., Boutrel, B., Chen, S. A., Kenny, P. J., Markou, A., O’Dell, L. E., Parsons, L. H. & Sanna, P. P. 2004. Neurobiological mechanisms in the transition from drug use to drug dependence. Neurosci Biobehav Rev, 27, 739–749.

Koob, G. F. & Volkow, N. D. 2010. Neurocircuitry of addiction. Neuropsychopharmacology, 35, 217–38.

Lagerspetz, K. & Portin, R. 1968. Simulation of cues eliciting aggressive responses in mice at two age levels. J Genet Psychol, 113, 53–63.

Leao, R. M., Cruz, F. C., Vendruscolo, L. F., De Guglielmo, G., Logrip, M. L., Planeta, C. S., Hope, B. T., Koob, G. F. & George, O. 2015. Chronic nicotine activates stress/reward-related brain regions and facilitates the transition to compulsive alcohol drinking. j Neurosci, 35, 6241–53.

Lubman, A., Emrick, C., Mosimann, W. F. & Freedman, R. 1983. Altered mood and norepinephrine metabolism following withdrawal from alcohol. Drug Alcohol Depend, 12, 3–13.

Macey, D. J., Schulteis, G., Heinrichs, S. C. & Koob, G. F. 1996. Time-dependent quantifiable withdrawal from ethanol in the rat: effect of method of dependence induction. Alcohol, 13, 163–170.

Marcinkiewcz, C. A., Dorrier, C. E., Lopez, A. J. & Kash, T. L. 2015. Ethanol induced adaptations in 5-HT2c receptor signaling in the bed nucleus of the stria terminalis: implications for anxiety during ethanol withdrawal. Neuropharmacology, 89, 157–167.

Martinotti, G., Di Nicola, M., Romanelli, R., Andreoli, S., Pozzi, G., Moroni, N. & Janiri, L. 2007. High and low dosage oxcarbazepine versus naltrexone for the prevention of relapse in alcohol-dependent patients. Hum Psychopharmacol, 22, 149–156.

Mcclintick, M. N. & Grant, K. A. 2016. Aggressive temperament predicts ethanol self-administration in late adolescent male and female rhesus macaques. Psychopharmacology (Berl), 233, 3965–3976.

Miyata, H., Hironaka, N., Takada, K., Miyasato, K., Nakamura, K. & Yanagita, T. 2008. Psychosocial withdrawal characteristics of nicotine compared with alcohol and caffeine. Ann NY Acad Sci, 1139, 458–465.

O’Dell, L. E., Roberts, A. J., Smith, R. T. & Koob, G. F. 2004. Enhanced alcohol self-administration after intermittent versus continuous alcohol vapor exposure. Alcohol Clin Exp Res, 28, 1676–1682.

Pandey, S. C., Roy, A. & Zhang, H. 2003. The decreased phosphorylation of cyclic adenosine monophosphate (cAMP) response element binding (CREB) protein in the central amygdala acts as a molecular substrate for anxiety related to ethanol withdrawal in rats. Alcohol Clin Exp Res, 27, 396–409.

Pandey, S. C., Sakharkar, A.J., Tang, L. & Zhang, H. 2015. Potential role of adolescent alcohol exposure-induced amygdaloid histone modifications in anxiety and alcohol intake during adulthood. Neurobiol Dis, 82, 607–619.

Pleil, K. E., Lowery-Gionta, E. G., Crowley, N. A., Li, C., Marcinkiewcz, C. A., Rose, J. H., Mccall, N. M., Maldonado-Devincci, A. M., Morrow, A. L., Jones, S. R. & Kash, T. L. 2015. Effects of chronic ethanol exposure on neuronal function in the prefrontal cortex and extended amygdala. Neuropharmacology, 99, 735–749.

Rassnick, S., Heinrichs, S. C., Britton, K. T. & Koob, G. F. 1993. Microinjection of a corticotropin-releasing factor antagonist into the central nucleus of the amygdala reverses anxiogenic-like effects of ethanol withdrawal. Brain Res, 605, 25–32.

Richardson, H. N., Lee, S. Y., O’Dell, L. E., Koob, G. F. & Rivier, C. L. 2008a. Alcohol self-administration acutely stimulates the hypothalamic-pituitary-adrenal axis, but alcohol dependence leads to a dampened neuroendocrine state. Eur J Neurosci, 28, 1641–53.

Richardson, H. N., Zhao, Y., Fekete, E. M., Funk, C. K., Wirsching, P., Janda, K. D., Zorrilla, E. P. & Koob, G. F. 2008b. MPZP: a novel small molecule corticotropin-releasing factor type 1 receptor (CRF1) antagonist. Pharmacol Biochem Behav, 88, 497–510.

Riihioja, P., Jaatinen, P., Haapalinna, A., Kiianmaa, K. & Hervonen, A. 1999. Effects of dexmedetomidine on rat locus coeruleus and ethanol withdrawal symptoms during intermittent ethanol exposure. Alcohol Clin Exp Res, 23, 432–438.

Riihioja, P., Jaatinen, P., Oksanen, H., Haapalinna, A., Heinonen, E. & Hervonen, A. 1997. Dexmedetomidine, diazepam, and propranolol in the treatment of ethanol withdrawal symptoms in the rat. Alcohol Clin Exp Res, 21, 804–808.

Riittinen, M. L., Lindroos, F., Kimanen, A., Pieninkeroinen, E., Pieninkeroinen, I., Sippola, J., Veilahti, J., Bergstrom, M. & Johansson, G. 1986. Impoverished rearing conditions increase stress-induced irritability in mice. Dev Psychobiol, 19, 105–111.

Roberts, A. J., Cole, M. & Koob, G. F. 1996. Intra-amygdala muscimol decreases operant ethanol self-administration in dependent rats. Alcohol Clin Exp Res, 20, 1289–1298.

Roberts, A. J., Heyser, C. J., Cole, M., Griffin, P.Koob, G. F. 2000. Excessive ethanol drinking following a history of dependence: animal model of allostasis. Neuropsychopharmacology, 22, 581–594.

Rogers, J., Wiener, S. G. & Bloom, F. E. 1979. Long-term ethanol administration methods for rats: advantages of inhalation over intubation or liquid diets. Behav Neural Biol, 27, 466–486.

Rylkova, D., Shah, H. P., Small, E. & Bruijnzeel, A. W. 2009. Deficit in brain reward function and acute and protracted anxiety-like behavior after discontinuation of a chronic alcohol liquid diet in rats. Psychopharmacology (Berl), 203, 629–640.

Sabino, V., Cottone, P., Koob, G. F., Steardo, L., Lee, M. J., Rice, K. C. & Zorrilla, E. P. 2006. Dissociation between opioid and CRF1 antagonist sensitive drinking in Sardinian alcohol-preferring rats. Psychopharmacology (Berl), 189, 175–186.

Schuckit, M. A., Tipp, J. E., Reich, T., Hesselbrock, V. M. & Bucholz, K. K. 1995. The histories of withdrawal convulsions and delirium tremens in 1648 alcohol dependent subjects. Addiction, 90, 1335–1347.

Schulteis, G., Markou, A., Cole, M.Koob, G. F. 1995. Decreased brain reward produced by ethanol withdrawal. Proc Natl Acad Sci USA, 92, 5880–5884.

Seif, T., Chang, S. J., Simms, J. A., Gibb, S. L., Dadgar, J., Chen, B. T., Harvey, B. K., Ron, D., Messing, R. O., Bonci, A. & Hopf, F. W.2013. Cortical activation of accumbens hyperpolarization-active NMDARs mediates aversion-resistant alcohol intake. Nat Neurosci, 16, 1094–1100.

Shibasaki, M., Kurokawa, K., Mizuno, K. & Ohkuma, S. 2012. Effect of aripiprazole on anxiety associated with ethanol physical dependence and on ethanol-induced place preference. J Pharmacol Sci, 118, 215–224.

Sommer, W. H., Rimondini, R., Hansson, A. C., Hipskind, P. A., Gehlert, D. R., Barr, C. S. & Heilig, M. A. 2008. Upregulation of voluntary alcohol intake, behavioral sensitivity to stress, and amygdala crhr1 expression following a history of dependence. Biol Psychiatry, 63, 139–45.

Staples, M. C., Kim, A. & Mandyam, C. D. 2015. Dendritic remodeling of hippocampal neurons is associated with altered NMDA receptor expression in alcohol dependent rats. Mol Cell Neurosci, 65, 153–62.

Thorsell, A., Johnson, J. & Heilig, M. 2007. Effect of the adenosine A2a receptor antagonist 3,7-dimethyl-propargylxanthine on anxiety-like and depression-like behavior and alcohol consumption in Wistar Rats. Alcohol Clin Exp Res, 31, 1302–1307.

Valdez, G. R., Roberts, A. J., Chan, K., Davis, H., Brennan, M., Zorrilla, E. P. & Koob, G. F. 2002a. Increased ethanol self-administration and anxiety-like behavior during acute ethanol withdrawal and protracted abstinence: regulation by corticotropin-releasing factor. Alcohol Clin Exp Res, 26, 1494–1501.

Valdez, G. R., Roberts, A. J., Chan, K., Davis, H., Brennan, M., Zorrilla, E. P. & Koob, G. F. 2002b. Increased ethanol self-administration and anxiety-like behavior during acute ethanol withdrawal and protracted abstinence: regulation by corticotropin-releasing factor. Alcohol Clin Exp Res, 26, 1494–1501.

Valdez, G. R., Sabino, V. & Koob, G. F. 2004. Increased anxiety-like behavior and ethanol self-administration in dependent rats: reversal via corticotropin-releasing factor-2 receptor activation. Alcohol Clin Exp Res, 28, 865–872.

Valdez, G. R., Zorrilla, E. P., Roberts, A. J. & Koob, G. F. 2003. Antagonism of corticotropin-releasing factor attenuates the enhanced responsiveness to stress observed during protracted ethanol abstinence. Alcohol, 29, 55–60.

Varlinskaya, E. I., Kim, E. U. & Spear, L. P. 2017. Chronic intermittent ethanol exposure during adolescence: Effects on stress-induced social alterations and social drinking in adulthood. Brain Res, 1654, 145–156.

Vendruscolo, L. F., Barbier, E., Schlosburg, J. E., Misra, K. K., Whitfield, T. W.JR., Logrip, M. L., Rivier, C., Repunte-Canonigo, V., Zorrilla, E. P., Sanna, P. P., Heilig, M.Koob, G. F. 2012. Corticosteroid-dependent plasticity mediates compulsive alcohol drinking in rats. J Neurosci, 32, 7563–7571.

Vendruscolo, L. F. & Roberts, A. J. 2014. Operant alcohol self-administration in dependent rats: focus on the vapor model. Alcohol, 48, 277–286.

Vetreno, R. P., Yaxley, R., Paniagua, B. & Crews, F. T. 2016. Diffusion tensor imaging reveals adolescent binge ethanol-induced brain structural integrity alterations in adult rats that correlate with behavioral dysfunction. Addict Biol, 21, 939–953.

Walker, B. M., Drimmer, D. A., Walker, J. L., Liu, T., Mathe, A. A. & Ehlers, C. L. 2010. Effects of prolonged ethanol vapor exposure on forced swim behavior, and neuropeptide Y and corticotropin-releasing factor levels in rat brains. Alcohol, 44, 487–493.

Walker, B. M., Rasmussen, D. D., Raskind, M. A. & Koob, G. F. 2008. alpha-lnoradrenergic receptor antagonism blocks dependence-induced increases in responding for ethanol. Alcohol, 42, 91–7.

Williams, A. M., Reis, D. J., Powell, A. S., Neira, L. J., Nealey, K. A., Ziegler, C. E., Kloss, N. D., Bilimoria, J. L., Smith, C. E. & Walker, B. M. 2012. The effect of intermittent alcohol vapor or pulsatile heroin on somatic and negative affective indices during spontaneous withdrawal in Wistar rats. Psychopharmacology (Berl), 223, 75–88.

Woldbye, D. P., Ulrichsen, J., Haugbol, S. & Bolwig, T. G. 2002. Ethanol withdrawal in rats is attenuated by intracerebroventricular administration of neuropeptide Y. Alcohol Alcohol, 37, 318–321.

Zhao, Y., Weiss, F. & Zorrilla, E. P. 2007. Remission and resurgence of anxiety-like behavior across protracted withdrawal stages in ethanol-dependent rats. Alcohol Clin Exp Res, 31, 1505–15.

Zorrilla, E. P., Heilig, M., De Wit, H. & Shaham, Y. 2013. Behavioral, biological, and chemical perspectives on targeting CRF(1) receptor antagonists to treat alcoholism. Drug Alcohol Depend, 128, 175–186.

